# Rtt107 cooperates with Rad55 or Slx4 to maintain genome stability in *Saccharomyces cerevisiae*

**DOI:** 10.1101/2024.02.05.579050

**Authors:** Joshua A. R. Brown, Michael S. Kobor

## Abstract

A range of genome maintenance factors respond to endogenous and exogenous DNA damage to prevent mutations and cell death. The scaffold protein, Rtt107, is important for the growth of cells exposed to DNA-damaging agents in the budding yeast *Saccharomyces cerevisiae*. Rtt107 binds to a diverse array of partner proteins, such as Slx4, and responds to DNA damage by localizing to phosphorylated histone H2A. Rad55–Rad57, a heterodimer involved in DNA repair, also binds to Rtt107, but the function of the Rtt107–Rad55–Rad57 complex remains unclear. In addition to their sensitivity to DNA-damaging agents, *rtt107*Δ mutants exhibit spontaneous genome instability phenotypes, including spontaneous loss of heterozygosity (LOH) caused by crossovers and other genetic events. However, the binding partners with which Rtt107 interacts to prevent spontaneous genome instability have yet to be elucidated. Here, we showed that Rtt107 acts in the same pathway as Rad55 to limit LOH, specifically by preventing crossover events. A *rad55-S404A* phosphorylation site mutation largely disrupted the interaction between Rtt107 and Rad55–Rad57, resulting in increased LOH and crossover rates, consistent with the contribution of Rtt107–Rad55–Rad57 interaction to genome stability. Strikingly, an *rtt107-K887M* mutation that reduces Rtt107 recruitment to H2A did not result in an LOH phenotype, suggesting that the role of Rtt107 in preventing LOH is distinct from its function as an H2A-binding scaffold. Rtt107 did not function primarily in the same pathway as Rad55 to limit recombination at the sensitive ribosomal DNA (rDNA) locus, but instead acted with Slx4 to maintain rDNA stability, suggesting that interactions of Rtt107 with different partners prevented distinct types of instability. Taken together, our observations suggested that Rtt107 limits spontaneous LOH and crossover events in part by binding to Rad55 in a manner dependent on Rad55-S404.

**Author Summary:** Numerous proteins are involved in the repair of damaged DNA and prevention of genome instability in cells, which would otherwise result in persistent changes to DNA. Genome maintenance pathways are evolutionarily conserved, and the budding yeast, *Saccharomyces cerevisiae*, is a powerful model organism for investigating the maintenance of genome integrity. Rtt107 is a scaffold protein expressed in yeast, containing conserved protein domains that are important for the function of genome maintenance proteins. Although the functions of Rtt107 in cells treated with DNA-damaging agents have been characterized in some detail, it remains unclear how Rtt107 prevents spontaneous genome instability in cells growing under normal conditions. Here, we found that Rtt107 prevents specific types of spontaneous genome instability and acts in the same pathway as the DNA repair protein, Rad55, to which it binds. Mutation of a possible Rtt107 binding site on Rad55 showed that Rtt107 indeed limited genome instability in part by binding to Rad55. Strikingly, Rtt107 also showed Rad55-independent roles in preventing ribosomal DNA (rDNA) instability, and in this context Rtt107 cooperated in part with its binding partner Slx4. Taken together, our results revealed the pathways by which the evolutionarily conserved protein Rtt107 limits spontaneous genome instability.

## 1 Introduction

The genome is constantly under threat by a variety of endogenous and exogenous factors, which are counteracted by the DNA damage response (DDR) consisting of interconnected pathways that detect, signal, and repair DNA damage [1,2]. These DDR proteins play crucial roles in ultimately preventing cell death when DNA damage occurs. Genome instability, encompassing both mutations and genomic rearrangements, can also occur as a byproduct of DNA damage and repair. Cells therefore rely on a variety of genome maintenance factors, with functions in the DDR and in processes such as DNA replication, to both survive DNA damage and limit genome instability.

DNA double-strand breaks (DSBs) are among the most deleterious types of DNA damage, and can cause cell death if left unrepaired [3]. DSBs and DSB repair can result in a range of types of genome instability, including deletions, gross chromosomal rearrangements (GCRs), and loss of heterozygosity (LOH) [2,3]. DSBs are repaired by a number of different pathways depending on the stage of the cell cycle at which they occur and the nature of specific lesions. In the budding yeast, *Saccharomyces cerevisiae*, DSB repair most frequently involves the homologous recombination (HR) pathway, while non-homologous end-joining (NHEJ) of DSB ends is specifically upregulated in G_1_ phase haploid yeast and is the predominant pathway in mammals [4]. HR occurs after resection, the 5′→3′ nucleolytic digestion of DSB ends that exposes single-stranded DNA (ssDNA) and leads to formation of a Rad51–ssDNA filament, and can generate reciprocal crossovers (RCOs) [3]. Alongside specific repair processes, DNA damage prompts a broader DDR that involves checkpoint signaling, which arrests the cell cycle while DNA damage is repaired [5]. Here, histone H2A-S129 phosphorylation (γH2A) is placed by the Mec1 and Tel1 sensor kinases around lesions, and forms a platform for the recruitment of other DDR factors, including checkpoint proteins [6]. Another histone mark, H4T80 phosphorylation, is placed near sites of DNA damage by Cla4, although the functional consequences of this chromatin mark in the DDR have yet to be fully elucidated [7].

Rtt107 is an evolutionarily conserved protein that plays multiple roles in the DDR, and is required for normal cell growth during chronic exposure to DNA-damaging agents [8]. At the molecular level, Rtt107 localizes to stalled replication forks, DSBs, and other DNA damage sites, where it acts as a scaffold to recruit other proteins to these lesions [9–14]. Rtt107 promotes recovery from treatments that damage DNA and the restart of DNA replication after damage repair, at least in part by limiting DNA damage and preventing checkpoint hyperactivation [8,15–17]. Rtt107 contains six BRCA1 C-terminal (BRCT) domains, which generally mediate both phosphorylation-dependent and -independent protein–protein interactions [8,18], and localizes near DSBs by binding γH2A via BRCT5/6 [10,19]. Rtt107 is also recruited near DSBs by binding phosphorylated H4T80 via BRCT1–4, with this representing the first identified molecular interaction partner of phosphorylated H4T80 [7]. The interaction of Rtt107 with γH2A is mediated by Rtt107-K887 and is required for its localization near DSBs [10,19], whereas interaction with phosphorylated H4T80 is mediated by Rtt107-K426 and partially contributes to its localization near DSBs [7]. Both *rtt107-K426A* and *rtt107-K887M* mutants exhibit hypersensitivity to chronic treatment with DNA-damaging agents, suggesting that Rtt107 confers DNA damage resistance, at least in part, by localizing to histone marks [10]. Rtt107–BRCT1–4 forms a tetrahedral structure that binds a variety of complexes, and in turn Rtt107 may act as a scaffold by recruiting these partners to γH2A-containing regions, to which Rtt107 itself is tethered via BRCT5/6 [9,14,15,20].

Rtt107 has been studied most extensively in the context of its localization to γH2A and its constitutive interaction with the Slx4 endonuclease [10,14,15]. DNA damage induces interaction of the Rtt107–Slx4 complex with the replication factor, Dpb11 [21]. In turn, this Rtt107– Slx4–Dpb11 interaction can competitively reduce the interaction between Dpb11 and the checkpoint factor, Rad9, while increased Rad9–Dpb11 interaction in the absence of the Rtt107–Slx4 complex can limit DSB resection and may contribute to checkpoint hyperactivation [16,17,22–24]. Slx4 is also required for phosphorylation of the SQ/TQ motifs of Rtt107 by Mec1 after treatment with the DNA-damaging agent, methyl methanesulfonate (MMS) [8,15]. However, it is unclear which functions of Rtt107 are dependent on its phosphorylation [8,21,25].

Rtt107 interacts with the Rad55–Rad57 heterodimer via Rtt107–BRCT1–4, and Rad55 contains an Rtt107 interaction motif that was shown to interact with Rtt107–BRCT1–4 *in vitro* [14,20]. However, the molecular function of the Rtt107–Rad55–Rad57 complex has yet to be determined. In general, Rad55–Rad57 acts as a recombination mediator, which promotes Rad51– ssDNA filament assembly for HR [26–30]. In contrast to the *rtt107*Δ mutant [8], both *rad55*Δ and *rad57*Δ mutants are sensitive to treatment with ionizing radiation [27,31], which induces DSBs. These findings suggest that Rtt107 may be less important for successful DSB repair than Rad55–Rad57, or alternatively that the importance of Rtt107 is limited to specific contexts. In addition to promoting DSB repair as a recombination mediator, Rad55–Rad57 also plays a role in DNA damage tolerance pathways that allow DNA lesions to be bypassed during replication and gap filling after replication [32]. Here, Rad55–Rad57 promotes the error-free postreplication repair (PRR) tolerance pathway, also known as template switching, by bridging Rad51 to the Shu complex [33–36]. Recent work also suggested that Rad55–Rad57 may downregulate the translesion synthesis (TLS) pathway [37]. Although *rtt107*Δ mutants exhibit spontaneous Rad52 foci [38], a marker of DNA damage and recombination, it has not been determined whether Rtt107 is important for any DNA damage repair function of Rad55–Rad57.

The function of Rtt107 in the context of exposure to DNA-damaging agents has been documented in detail, while the role of Rtt107 in preventing spontaneous genome instability is only beginning to be elucidated. Originally, the *rtt107*Δ mutant was identified in screens for mutants with elevated levels of spontaneous LOH and GCRs, chromosomal instability, and phenotypes reflecting spontaneous DNA damage foci and DDR signaling [38–42]. The *rtt107*Δ mutant also exhibits hyperrecombination at the ribosomal DNA (rDNA) locus [38], a highly transcribed series of tandem repeats spanning ∼10% of the *S. cerevisiae* genome that represents a unique challenge to the maintenance of genome stability [43]. Taken together, these phenotypes indicate that Rtt107 plays an important role in preventing several types of spontaneous genome instability. However, it remains to be explored whether the role of Rtt107 in preventing spontaneous instability is related to its established functions in the response to exogenous DNA-damaging agents. Rtt107 binds to a variety of DDR complexes in addition to Slx4 and Rad55– Rad57, and it is possible that Rtt107 cooperates with one or several of its documented binding partners to prevent spontaneous genome instability. These partners include the SMC5/6 (Structural Maintenance of Chromosomes) complex, Rtt101–Mms1–Mms22, and Mus81– Mms4–Cdc7–Dbf4–Cdc5 [9,12,21,44–46]. Further, Rtt107 was shown to interact with the Sgs1 helicase in a yeast two-hybrid screen [20].

Genetic interactions underlying genome instability phenotypes represent powerful tools for investigating gene function, and epistatic interactions can often be interpreted in the framework of classical biochemical pathway models [2]. Here, we made use of a comprehensive suite of genetic analyses to uncover the specific pathways in which Rtt107 prevents specific types of spontaneous genome instability. First, we established that the function of Rtt107 in preventing spontaneous LOH was distinct from its role as a γH2A-binding scaffold and from the role of Rtt107 in the context of chronic exposure to DNA-damaging agents, but that Rtt107– BRCT5/6 was still important for preventing spontaneous LOH. Next, we found that Rtt107 cooperated specifically with Rad55 to limit LOH as well as mitotic RCO events. A *rad55-S404A* mutant showed reduced interaction between Rtt107 and Rad55 and increased rates of mitotic RCOs and LOH, although the *rtt107*Δ and *rad55-S404A* mutations also made independent contributions to the LOH rate. However, rather than functioning with Rad55 to limit spontaneous recombination at the rDNA locus, Rtt107 cooperated with Slx4 to maintain rDNA stability. Taken together, these results demonstrated that the role of Rtt107 in preventing LOH was largely independent of its previously identified function as a γH2A-binding scaffold, and indicated that Rtt107 cooperated with Rad55 or Slx4 to prevent different types of instability at distinct genetic loci.

## 2 Materials and Methods

### 2.1 Yeast strains and plasmids

Yeast strains used in this study are listed in Table S1, made using standard genetic techniques. The primers used to generate the Rad57-HA construct also encoded a GCAGCAGCAGCAGCAGCAGCA linker between Rad57 and the HA tag (primers available upon request).

The plasmids used in this study are listed in Table S2, and were produced using standard genetic techniques. Plasmids pMK429 and pMK430 were generous gifts from Dr. Jocelyn Krebs (University of Alaska Anchorage), and were used in the experiments presented in Figure 2C to examine the *H2A-S129A* allele.

### 2.2 Genome instability assays

Several assays for genome instability phenotypes were applied in this study under normal growth conditions. Each assay is briefly summarized below and outlined schematically in Fig. 1.

**Fig. 1.**
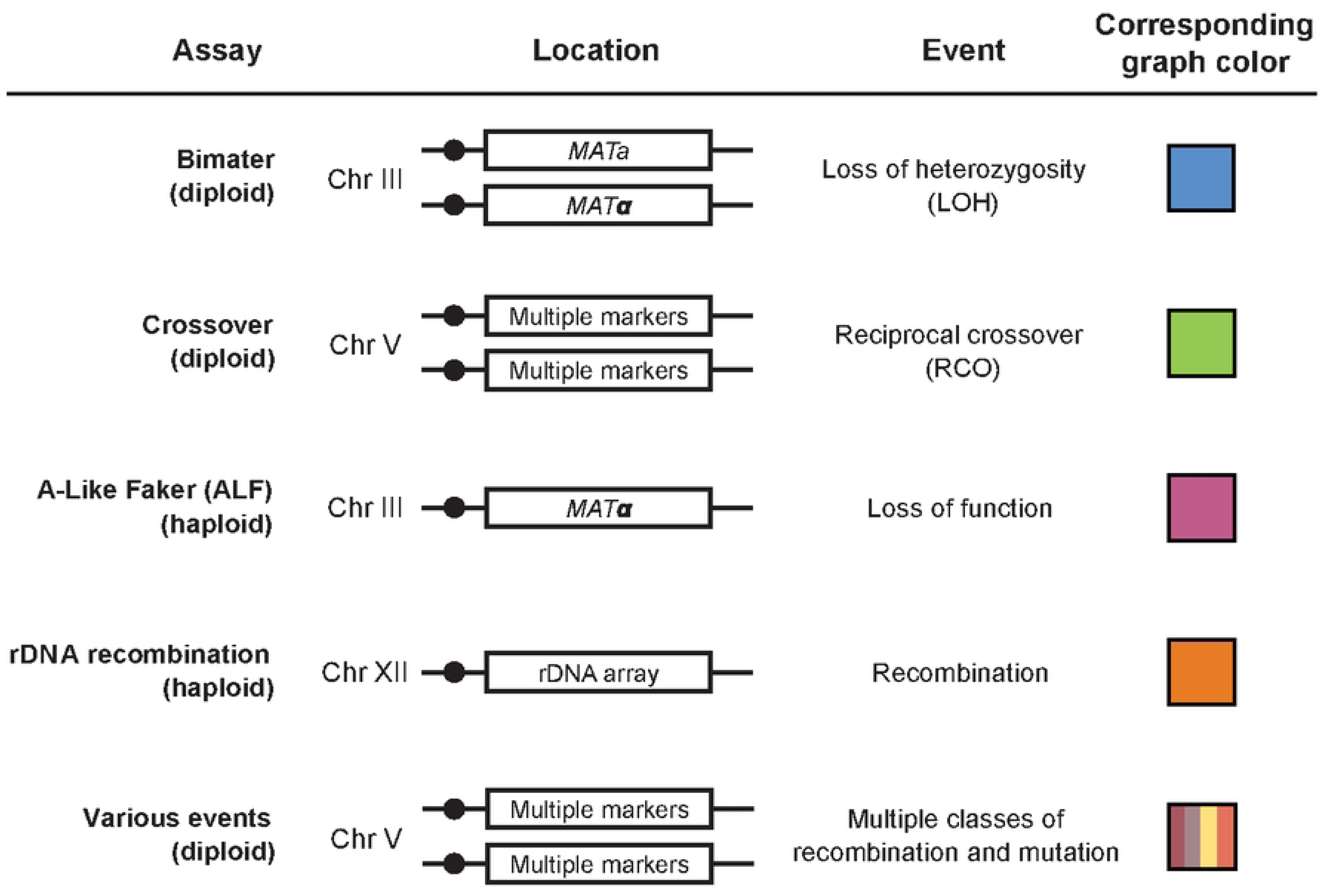
Targeted spontaneous genome instability assays used in this study. The locations of mutations and genomic rearrangements in each assay are shown. All assays were performed under normal growth conditions. The graph colors shown on the right correspond to the colors of graphs for each assay in subsequent figures.

#### 2.2.1 Quantitative bimater assay

The procedure for the quantitative bimater assay was adapted from a previous report [25]. An overnight culture of a *MAT*a mating tester strain was grown in YPD at 30°C, diluted, and plated onto solid medium containing no amino acids (−AA plates). Overnight cultures of the test strain were grown at 30°C and diluted. Cultures were then plated at different dilutions onto −AA plates to count mating events with the *MAT*a tester and onto YPD plates to score total counts. Plates were grown at 30°C for 2–3 days before counting. Mating rates and 95% confidence intervals were calculated with the maximum likelihood estimate method, implemented using the `newton.LD.plating` function in the `rSalvador` R package [47]. Statistical significance was assessed by comparing the overlap of confidence intervals [48], and comparisons between strains were considered significant if the 95% confidence intervals did not overlap. This approach is more stringent than a conventional *p*-value cutoff of 0.05 [48].

The bimater assay shown in Figure 2B was adapted to maintain plasmid selection. Strains were grown in SC-Leu medium, and mated to a *leu2*Δ mating tester strain.

**Fig. 2.**
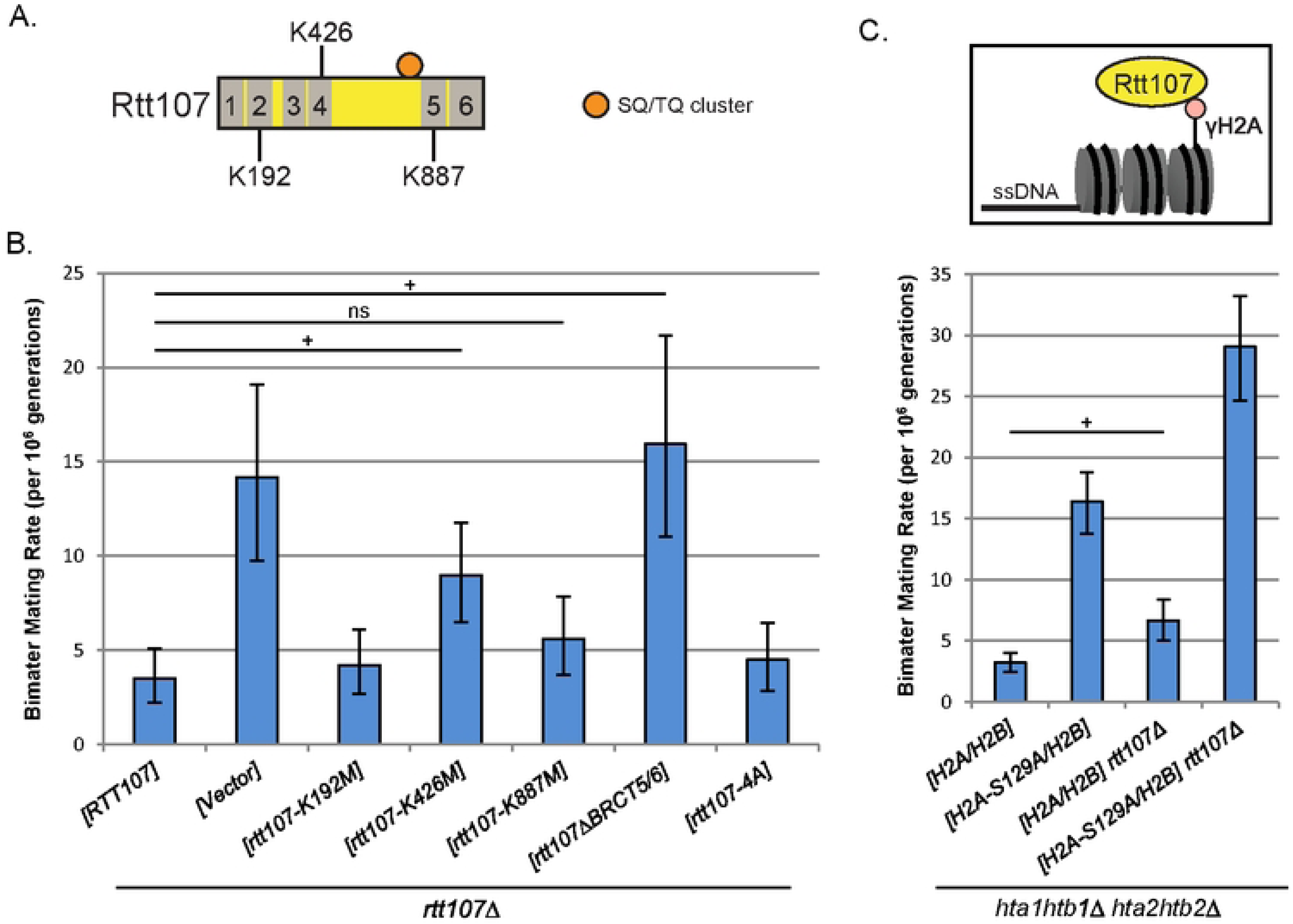
Rtt107–BRCT1–4 and Rtt107–BRCT5/6 were important for limiting spontaneous LOH. A) Rtt107 structural features mutated in panel B. BRCT domains are numbered and shown in gray. Target lysines are labeled in the center of the BRCT domain in which they are located. The orange circle indicates the SQ/TQ cluster of Rtt107, targeted by the *rtt107-4A* mutation. B) Rtt107-K426 and Rtt107–BRCT5/6 contributed to genome maintenance in the bimater assay, performed using homozygous diploid strains. Mating rates and 95% confidence intervals were calculated using the maximum likelihood method, and statistical significance was assessed by the non-overlap of confidence intervals (see *Materials and Methods*). +, non-overlap of 95% confidence intervals between two strains (statistically significant); ns, overlap of confidence intervals (not significant). Strains indicated in this panel were in the W303 *RAD5* background, and were grown in SC-Leu. At least eight independent colonies were tested. C) Rtt107 prevented spontaneous LOH independently of γH2A in the bimater assay. The *H2A-S129A* mutation caused genome instability, and the *rtt107*Δ *H2A-S129A* double mutant phenotype was greater than those of either single mutant. Mating rates and 95% confidence intervals were calculated and compared as described for panel B. Eleven independent colonies were tested.

#### 2.2.2 Reciprocal crossover (RCO) assay

RCO events were examined using *can1-100* and *SUP4-o* alleles on chromosome V, as described previously [49]. Independent colonies were grown on YPD plates for 2 days and resuspended in water. Cells were plated at different dilutions onto solid SC-Arg medium with or without 250 μg/mL of canavanine sulfate (Sigma). Plates were grown for 4 days at 30°C and stored at 4°C for 1 day to allow colony color development. Red/white sectored colonies on plates containing canavanine were scored as representative of RCO events in the interval between *CEN5* and *CAN1* loci. Counts were compared to colony counts on plates lacking canavanine, and the frequency of RCOs was calculated. Statistical significance was assessed using Student’s *t* test.

#### 2.2.3 Quantitative A-like faker (ALF) assay

A quantitative version of the A-like faker (ALF) assay was performed using the same method as the bimater assay, except that *MATα* haploid strains were examined for their ability to mate with a *MATα* tester strain. Mating rates and statistical significance were calculated using the same methods as described above.

#### 2.2.4 rDNA recombination assay

An rDNA recombination assay was adapted from a previous report to allow for quantification by fluctuation analysis [50]. Strains were plated onto solid medium lacking adenine and grown for 2 days at 30°C. Independent cultures were grown in liquid medium at 30°C for 2 days. Diluted cultures were plated onto solid medium lacking arginine with or without 120 μg/mL canavanine (Sigma). Plates with canavanine were incubated for 2 days, replica plated onto media with or without adenine, and incubated for 2 days. Canavanine-resistant colonies that did not grow on plates lacking adenine were scored as representative of rDNA recombination. Scored colonies were compared to colony counts from plates lacking canavanine. The rDNA recombination rates and statistical significance were calculated by the method described for mating rates in the bimater assay.

#### 2.2.5 Diploid instability assay

The assay used to examine various types of genome instability in diploids shown in Figure 6C was adapted from a previous report [49]. Colonies were grown, diluted, and plated as described for the RCO assay. Colonies were plated at separate dilutions to examine Class 5/6 events due to their lower rates. After 4 days of growth at 30°C, colonies were replica plated onto several types of solid media to score growth phenotypes, as described previously [49]. Plates were grown at 30°C for 2 days and stored at 4°C for 2 days to allow colony color development. Non-sectored colonies were scored as representative of Class 1–6 events according to their respective growth phenotypes. Mutation rates and statistical significance were calculated as described for mating rates in the bimater assay. A total of 28 Class 3 mutants were examined by PCR as described previously [49] to discriminate between local gene conversion and *de novo* mutation events at the *SUP4-o* locus.

### 2.3 DNA damage sensitivity assays

Overnight cultures grown in YPD at 30°C were diluted to 0.5 OD_600_. Cells were then 10-fold serially diluted, and plated on YPD medium containing the indicated concentration of MMS, HU, or CPT (Sigma). Plates were incubated for 2 days at 30°C.

### 2.4 TCA extraction

Overnight cultures were diluted to 0.3 OD_600_ in YPD medium and grown to mid-log phase. Whole-cell extracts were prepared by glass bead lysis in the presence of trichloroacetic acid (TCA). Samples were boiled for 5 min and resolved by 8% SDS-PAGE. Western blotting analysis was performed using anti-FLAG M2 (Sigma) and anti-PGK1 (Thermo Fisher) antibodies.

### 2.5 Analytical-scale protein–protein interaction assay

Overnight cultures were diluted to 0.3 OD_600_ in 4 L of YPD medium, and grown to mid-log phase. The procedure for co-immunoprecipitation was adapted from a previous report [51]. Briefly, cells were disrupted with a coffee grinder in the presence of dry ice pellets and resuspended in TAP-IP buffer (50 mM Tris, pH 7.8, 150 mM NaCl, 1.5 mM MgAc, 0.15% Nonidet P-40, 1 mM DTT, 10 mM NaPPi, 5 mM EGTA, 5 mM EDTA, 0.1 mM Na_3_VO_4_, 5 mM NaF, and Roche cOmplete™ Protease Inhibitor Mix). Crude extracts were prepared by centrifugation, and incubated for 2 hours at 4°C with anti-FLAG M2 antibody (Sigma)-coupled Dynabeads (Invitrogen). Beads were washed with TAP-IP buffer. Captured protein was detected by immunoblotting with anti-FLAG M2 antibody (Sigma) and visualized using an Odyssey Infrared Imaging System (Licor). Co-purifying proteins were imaged with anti-HA (Sigma) and anti-VSV (Bethyl Laboratories) antibodies. Anti-PGK1 (Thermo Fisher) and anti-VSV (Bethyl Laboratories) antibodies were applied to the input fraction as loading controls.

## 3 Results

### 3.1 Rtt107–BRCT1–4 and BRCT5/6 were important for limiting spontaneous LOH

Despite the identification of a number of Rtt107 binding partners and of the molecular functions of Rtt107 in cells exposed to DNA-damaging agents, it remains an important question how Rtt107 operates to prevent a variety of types of spontaneous genome instability. As Rtt107 acts as a scaffold by binding to different partners via Rtt107–BRCT1–4 and BRCT5/6, we first examined which structural features of Rtt107 were required for limiting spontaneous genome instability. For this purpose, we examined a spontaneous LOH phenotype that was previously identified for the *rtt107*Δ mutant using the bimater assay (Fig. 1) [25,39]; this assay determines the rate of LOH at the mating type locus in diploid yeast and therefore captures a range of underlying mechanisms, encompassing mitotic recombination, gene conversion, chromosome loss, and chromosomal rearrangements including deletions. We targeted previously studied Rtt107 features with point mutations and a C-terminal truncation (Fig. 2A), and examined the impacts of these mutations on spontaneous LOH rates. The interaction of Rtt107 with γH2A is largely disrupted by the *rtt107-K887M* mutation [10,19], and C-terminal truncation of Rtt107 after aa820 (*rtt107*Δ*BRCT5/*6) was used to disrupt any other protein–protein interactions that may require BRCT5/6 [10]. We used the *rtt107-K426M* mutation within BRCT1–4 to disrupt the binding of Rtt107 to phosphorylated H4T80 [7], although it should be noted that Rtt107– BRCT1–4 also has additional binding partners that could be affected by this specific point mutation [9,14,15,20]. We examined other regions of Rtt107 using nonphosphorylatable mutations of each SQ/TQ motif in the SQ/TQ cluster of Rtt107 (*rtt107-4A*) [8], and a mutation designed to disrupt the functions of Rtt107– BRCT1/2 (*rtt107-K192M*) [10]. First, we confirmed that plasmid-borne *RTT107* reduced the rate of LOH in the *rtt107*Δ mutant to close to that seen in wild-type cells (Fig. 2B) consistent with previous reports of a bimater phenotype for the *rtt107*Δ mutant [24,28]. Note that in these experiments, comparisons were considered statistically significant if the 95% confidence intervals did not overlap (see *Materials and Methods*). The *rtt107-K887M* mutation did not result in an increase in rate of LOH, suggesting that the function of Rtt107 in this context may be independent of its localization to γH2A. In contrast, the *rtt107*Δ*BRCT5/6* mutation resulted in an elevated LOH rate, similar to complete loss of Rtt107. This suggested that BRCT5/6 played a γH2A-independent role in genome maintenance. The *rtt107-K426M* mutation increased the rate of LOH in this assay, indicating that BRCT1–4 also contributed to LOH prevention. In contrast, the *rtt107-K192M* and *rtt107-4A* mutations did not affect rescue of the bimater phenotype. This was perhaps unsurprising in the case of the *rtt107-K192M* mutation, which has been shown not to disrupt Rtt107 function in other contexts [10]. Taken together, these data indicated that both Rtt107–BRCT1–4 and Rtt107– BRCT5/6 contributed to LOH prevention.

We next confirmed that the function of Rtt107 in this context was indeed independent of its localization to γH2A, as our data from the *rtt107-K887M* mutant suggested. For this purpose, we examined whether a nonphosphorylatable *H2A-S129A* mutation affected the ability of Rtt107 to limit spontaneous LOH. The results indicated that incorporation of the *H2A-S129A* allele increased the rate of LOH in the bimater assay, consistent with the function of γH2A in the DDR (Fig. 2C) [52]. The *rtt107*Δ mutation increased the rate of LOH of the *H2A-S129A* mutant indicating that Rtt107 and γH2A had independent functions in this context, and that Rtt107 did not prevent spontaneous LOH by acting as a scaffold localized to γH2A.

### 3.2 Rtt107 acted in the same pathway as Rad55 to limit spontaneous LOH

Having found that multiple domains of Rtt107 contributed to its function in maintaining genome stability but that Rtt107 localization to γH2A was not required, we next examined whether Rtt107 limited genome instability in concert with its known physical interaction partners. We first tested whether the role of Rtt107 in limiting spontaneous genome instability was related to its well-characterized function in downregulating the Rad9–Dpb11 interaction, which has been examined previously in the context of treatment with DNA-damaging agents [16]. To determine whether the genome instability of the *rtt107Δ* mutant was due to increased Rad9–Dpb11 interaction, we utilized a *rad9-ST462,474AA* allele (hereafter referred to as *rad9-2A*), which disrupts the Rad9–Dpb11 interaction and partially suppresses the MMS sensitivity of the *rtt107*Δ mutant [17,53]. We compared the *rtt107*Δ, *rad9-2A*, and *rtt107*Δ *rad9-2A* mutant phenotypes in the bimater assay, and found that the *rad9-2A* allele modestly increased the LOH rate compared to the wild-type control (Fig. S1). Introduction of the *rad9-2A* allele into the *rtt107*Δ mutant background increased the rate of LOH to approximately double that of the *rtt107*Δ single mutant, suggesting that spontaneous LOH in the *rtt107*Δ mutant was not caused by Rad9–Dpb11 interaction. It should be noted that strains carrying genomic *H2A/H2B* gene derivatives were used in this assay and subsequent experiments, and the *rtt107*Δ mutation caused a somewhat stronger phenotype in this context compared to strains with plasmid-borne *H2A/H2B* genes (Fig. 2C).

As Rtt107 did not prevent spontaneous LOH by regulating the Rad9–Dpb11 interaction or localizing to γH2A, we next examined whether Rtt107 limits genome instability by cooperating with its other known or putative physical interaction partners, specifically Slx4, SMC5/6, Mms22, Mms4, Sgs1, and Rad55 [9,12,21,44–46]. For this purpose, we compared the bimater phenotype of the *rtt107*Δ mutant to strains with mutations of Rtt107 physical interaction partners as well as corresponding double mutants (Fig. 3A–F). Specifically, we sought to identify a mutation in an Rtt107 interaction partner that had a similar spontaneous LOH phenotype to the *rtt107*Δ mutant, and that exhibited epistasis with the *rtt107*Δ mutation. We used a hypomorphic *smc6-9* allele to examine the essential SMC5/6 complex (Fig. 3B) [54], as the *smc6-9* mutant shows a defect similar to the *rtt107*Δ mutant in DSB relocalization to the nuclear periphery [55]. We used an *mms4-8A* allele to reduce the interaction between Rtt107 and Mus81–Mms4 (Fig. 3D) [45]. Of the mutations examined, only the *rtt107*Δ *rad55*Δ double mutant had a phenotype similar to those of the *rtt107*Δ and *rad55*Δ single mutants, indicating an epistatic interaction between the two mutations (Fig. 3F). This pattern was not observed for mutants of other Rtt107 binding partners, and was consistent with an overlap in function between Rtt107 and Rad55 in preventing spontaneous LOH.

**Fig. 3.**
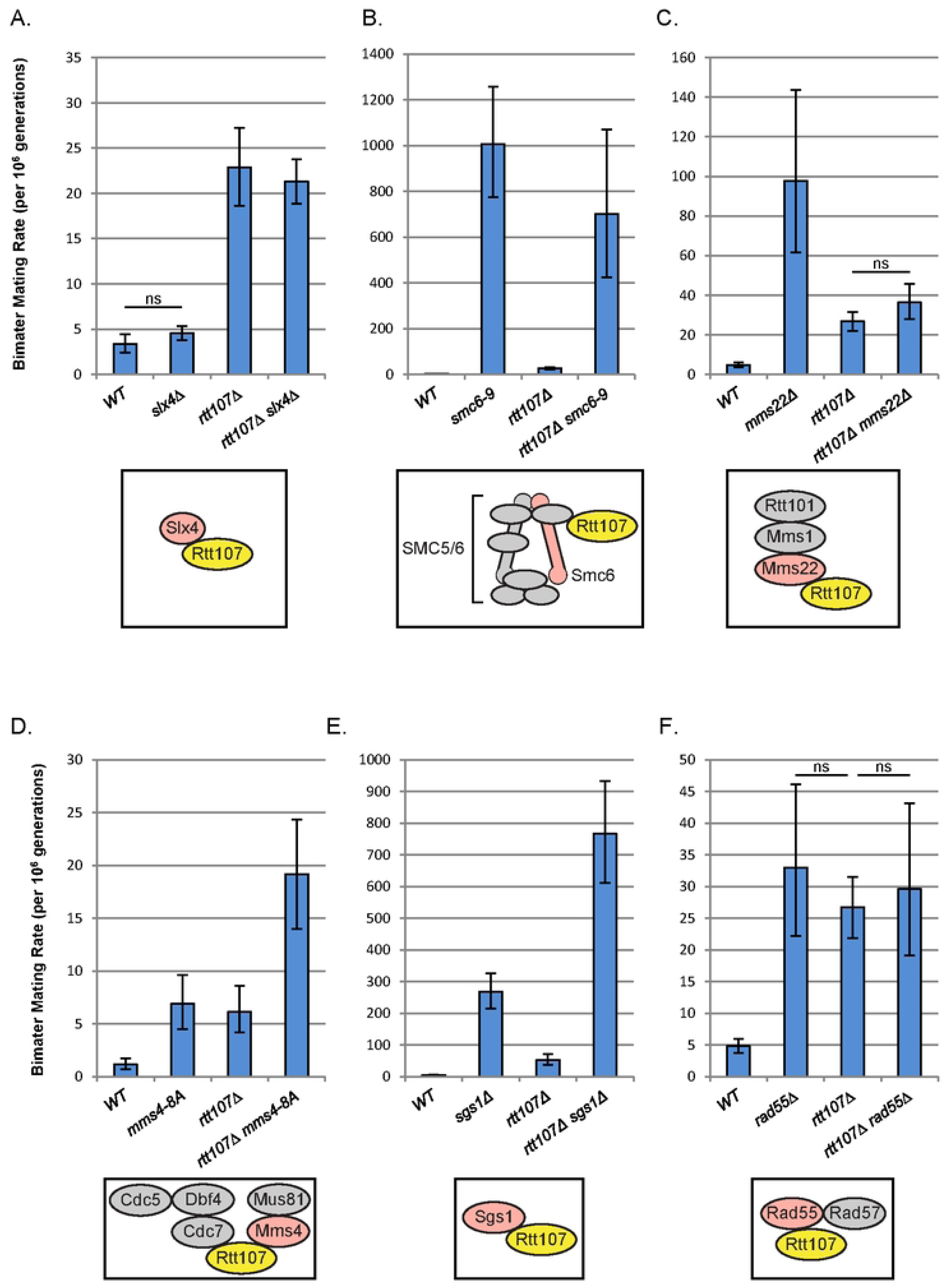
Rtt107 limited spontaneous LOH in the same pathway as Rad55. All panels depict bimater assays. The mating rates and 95% confidence intervals were calculated and compared as described for the bimater assay in Figure 2B. ns, overlap of confidence intervals (not significant). A simplified model of the corresponding physical interaction of Rtt107 is indicated beneath each assay, and mutated complex members are shown in color. Experiments were performed in the W303 *rad5-535* background except where noted. Note that the scale on the *y*-axis differs between panels. A) The *slx4*Δ mutation did not increase the rate of spontaneous LOH or affect the *rtt107*Δ mutant phenotype. At least six independent colonies were tested. B) The *smc6-9* mutation caused far greater instability in the bimater assay than the *rtt107*Δ mutant. The phenotype of the *rtt107*Δ *smc6-9* double mutant was not distinguishable from that of the *smc6-9* single mutant. At least seven independent colonies were tested. C) The *mms22*Δ mutation caused instability in the bimater assay. The *mms22*Δ mutant exhibited a higher rate of LOH than the *rtt107*Δ mutant, and this phenotype was suppressed by the *rtt107*Δ mutation. Ten to thirteen independent colonies were tested. D) The *rtt107*Δ *mms4-8A* double mutant exhibited a greater rate of LOH than either single mutant in the bimater assay. At least seven independent colonies were tested. Note that this experiment was performed in the W303 *RAD5* background, and that the complex model depicted in this panel is a hypothetical arrangement. E) The *sgs1*Δ mutant exhibited a higher LOH rate than the *rtt107*Δ mutant in the bimater assay. The *rtt107*Δ *sgs1*Δ double mutant exhibited a higher LOH rate than either single mutant. Eleven independent colonies were tested. F) The *rtt107*Δ *rad55*Δ double mutant exhibited a similar LOH rate to the *rtt107*Δ and *rad55*Δ single mutants. At least 12 independent colonies were tested.

### 3.3 Rtt107 acted in the same pathway as Rad55 to limit spontaneous mitotic RCOs

As the *rtt107*Δ, *rad55*Δ, and *rtt107*Δ *rad55*Δ mutants all exhibited a similar LOH phenotype, we next interrogated the relation between Rtt107 and Rad55 in greater detail. Given the complexities and interrelations between genes involved in the DDR, we first tested whether the relation observed between the *rtt107*Δ and *rad55*Δ mutations was dependent on strain background. Specifically, a *rad5-535* allele is present in the canonical W303 yeast strain background used here, and this can possibly affect findings of DDR studies [56]. To test whether this mutation might affect our results, we repeated the bimater assay in a *RAD5*-corrected W303 background. Reassuringly, we observed a similar pattern of LOH phenotypes (Fig. 4A), although perhaps not surprisingly the LOH rates for each strain were much lower than those in the *rad5-535* background (Fig. 3F). This indicated that the epistatic interaction between the *rtt107*Δ and *rad55*Δ mutations was not dependent on the presence of the *rad5-535* allele, although LOH rates may be mediated in some way by Rad5, and again suggested that Rtt107 may act in the same pathway as Rad55 to limit spontaneous LOH As a shared function for Rtt107 and Rad55 has not yet been identified, we sought to characterize the degree of cooperation between Rtt107 and Rad55 in limiting spontaneous genome instability in more detail. LOH in the bimater assay is predominantly mediated by crossover events produced by mitotic recombination in wild-type yeast cells [39]. Given that the Rad55–Rad57 heterodimer acts as a mediator of homologous recombination, which can produce mitotic RCO, we hypothesized that spontaneous RCO may occur in the *rtt107*Δ and *rad55*Δ mutants. Therefore, we tested whether Rtt107 and Rad55 limited the rate of spontaneous mitotic RCO, and whether they did so by acting within the same pathway. We examined the *rtt107*Δ and *rad55*Δ mutants by sectoring assay for spontaneous mitotic RCOs within a 120-kb interval on chromosome V in diploid cells, as described previously [49]. It should be noted that this and all subsequent experiments regarding the relation between *RTT107* and *RAD55* were performed in the *RAD5*-corrected W303 background, as described above. The *rtt107*Δ and *rad55*Δ mutants showed higher frequencies of spontaneous RCOs than the wild-type control (Fig. 4B). The *rtt107*Δ single mutant and *rtt107*Δ *rad55*Δ double mutant showed similar increases in mitotic RCO frequency, suggesting that Rtt107 and Rad55 limited RCOs by acting within the same pathway.

**Fig. 4.**
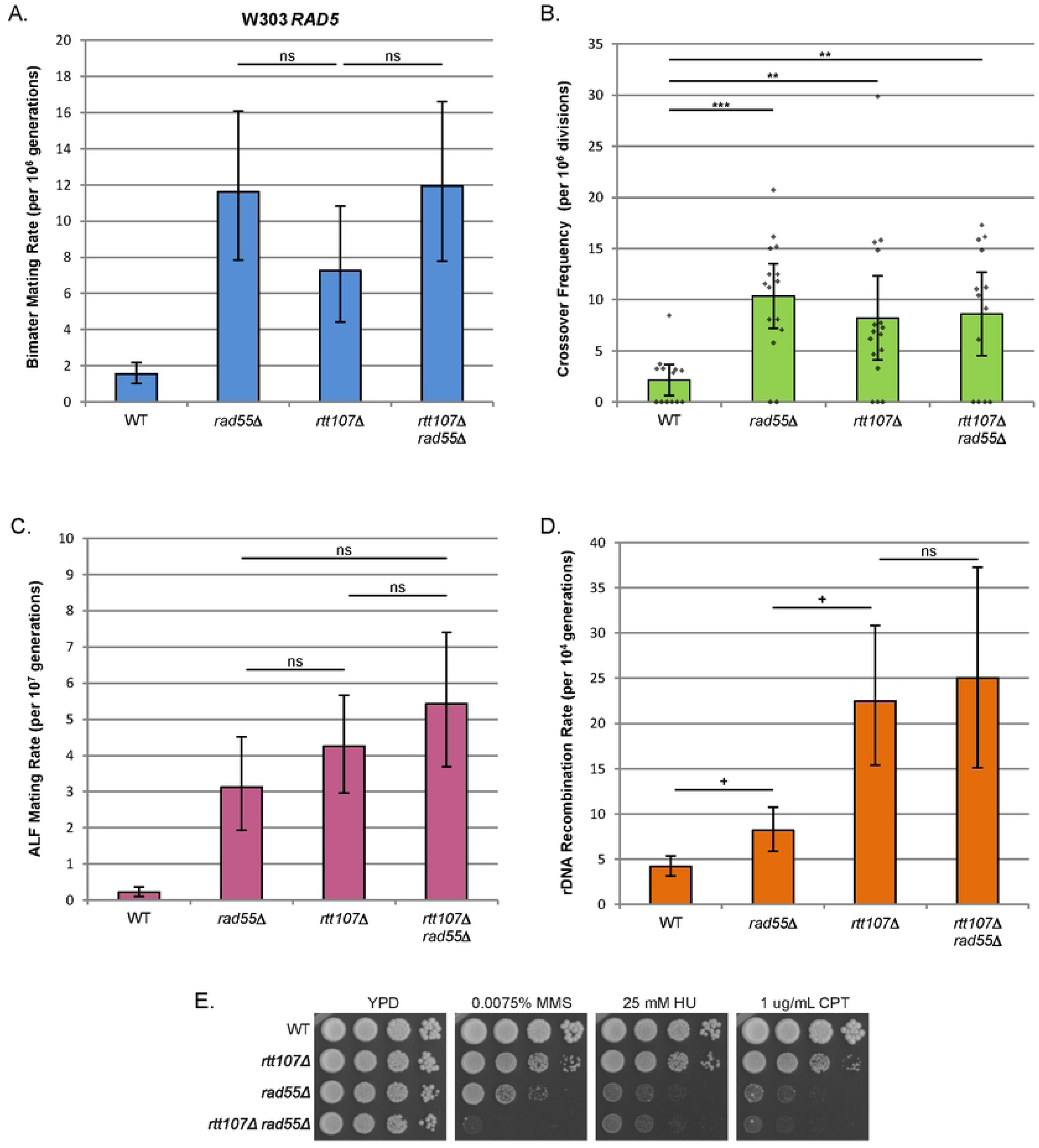
Rtt107 and Rad55 limited spontaneous mitotic RCO frequency in the same pathway. Experiments were performed in the W303 *RAD5* background. Rates and 95% confidence intervals for the bimater assay (panel A), ALF assay (panel C), and rDNA instability assay (panel D) were calculated and compared as described for the bimater assay in Figure 2B. +, non-overlap of 95% confidence intervals between two strains (statistically significant); ns, overlap of confidence intervals (not significant). A) The *rtt107*Δ *rad55*Δ double mutant exhibited a similar LOH rate to the *rad55*Δ single mutant in the bimater assay. Findings were similar to those in the W303 *rad5-535* background (Fig. 3F). Eight to 10 independent colonies were tested. B) The *rtt107*Δ, *rad55*Δ, and *rtt107*Δ *rad55*Δ mutants exhibited a similar increase in spontaneous mitotic RCO frequency. Canavanine-resistant sectored colonies were scored as representative of RCOs. Frequencies were calculated from 13–16 independent colonies. **, *p* < 0.01, *** *p* < 0.005 compared to the wild-type (Student’s *t* test). C) The *rtt107*Δ and *rad55*Δ mutants exhibited a higher rate of instability in the haploid ALF assay. The phenotype of the *rtt107*Δ *rad55*Δ double mutant showed a trend toward a higher rate than the *rtt107*Δ mutant, but this difference was not statistically significant. Haploid strains were used to determine mating rates in the ALF assay, and 7–9 independent colonies were tested. D) The *rtt107*Δ and *rad55*Δ mutants exhibited a higher rate of spontaneous rDNA recombination in comparison to wild-type. The *rtt107*Δ mutant and *rtt107*Δ *rad55*Δ double mutant showed a higher rate of rDNA recombination than the *rad55*Δ mutant. rDNA recombination was measured as the loss of an *ADE2-CAN1* marker in the rDNA, and eight independent colonies were tested. E) The *rtt107*Δ *rad55*Δ double mutant showed greater sensitivity to MMS treatment than either single mutant. Tenfold serial dilutions from overnight cultures were plated on media containing the indicated concentrations of DNA-damaging agents, and then incubated for 2 days at 30°C.

Next, we tested whether Rtt107 and Rad55 cooperated to limit other genomic events in addition to RCOs that may have caused LOH in the diploid bimater assay, such as gene conversion, chromosomal rearrangements, and chromosome loss. Each of these events, other than RCOs, can be captured by the ALF assay that measures loss of mating locus function in haploid cells [39]. Consistent with an earlier screen, quantitative ALF assay showed that the *rtt107*Δ and *rad55*Δ mutants both exhibited higher mating rates than the wild-type (Fig. 4C) [39]. The *rtt107*Δ *rad55*Δ double mutant phenotype trended toward a slightly higher mating rate than either of the single mutants, although the difference was not statistically significant. These observations suggested that Rtt107 and Rad55 may partly act within the same pathway to limit the types of genome instability that are captured by the ALF assay. However, as it remained formally possible that Rtt107 and Rad55 had independent functions in this context, we focused on exploring other genome instability phenotypes.

The observations outlined above suggested that Rtt107 cooperated with Rad55 to prevent specific types of genome instability at the mating locus and on chromosome V in diploid cells. Next, to test whether Rtt107 prevented various types of spontaneous instability more broadly at different genetic loci, we examined spontaneous recombination at the sensitive rDNA locus [43]. Specifically, we used a quantitative version of an assay for rDNA recombination based on the loss of an *ADE2*-*CAN1* marker in the rDNA of haploid yeast to monitor rDNA instability [50]. The *rtt107*Δ mutant exhibited rDNA hyperrecombination, as reported previously (Fig. 4D) [38]. The *rad55*Δ mutant exhibited a modest (∼2-fold) increase in spontaneous rDNA recombination, which was significantly less than that in the *rtt107*Δ mutant (∼5-fold), suggesting that Rtt107 did not primarily maintain rDNA stability by cooperating with Rad55. Consistent with largely separate roles of Rtt107 and Rad55 in rDNA maintenance, the *rtt107*Δ *rad55*Δ double mutant showed a trend toward greater instability than the *rtt107*Δ single mutant, although this increase was not statistically significant. We considered that Rtt107 may instead maintain rDNA stability with its major binding partner, Slx4, which maintains rDNA structure during DNA replication [57]. Both the *slx4*Δ single mutant and *rtt107*Δ *slx4*Δ double mutant exhibited rDNA hyperrecombination at rates similar to that of the *rtt107*Δ mutant (Fig. S2), which suggested that Rtt107 maintained rDNA stability by cooperating with Slx4 rather than Rad55.

Our results up to this point supported substantial overlap in function between Rtt107 and Rad55 in preventing specific types of spontaneous genome instability. We next examined the degree of cooperation between Rtt107 and Rad55 during chronic treatment with DNA-damaging agents to clarify the degree of cooperation between Rtt107 and Rad55 in the DDR. Focusing on agents previously shown to cause sensitivity to strains lacking Rtt107, we examined whether Rtt107 and Rad55 acted in the same DDR pathway to confer resistance to the DNA-damaging agents MMS, hydroxyurea (HU), or camptothecin (CPT) in a growth assay (Fig. 4E). Given the extreme sensitivity of the *rad55*Δ mutant to these agents, phenotypic comparison of the *rad55*Δ mutant and *rtt107*Δ *rad55*Δ double mutant required their use at low concentrations. However, it should be noted that the reported sensitivity of the *rtt107*Δ mutant to these agents was not readily apparent at the low concentrations used in this study [8,58]. We first confirmed that the *rtt107*Δ *rad55*Δ mutant exhibited enhanced MMS sensitivity compared to single mutants, as reported previously [20]. Next, we found that the *rad55*Δ mutant exhibited much greater sensitivity to HU and CPT than the *rtt107*Δ mutant. The *rtt107*Δ *rad55Δ* double mutant also showed similar HU sensitivity and a slight but reproducible increase in CPT sensitivity compared to the *rad55*Δ single mutant (Fig. 4E). Taken together, these results suggested that Rtt107 was not required for the function of Rad55 in the context of resistance to chronic DNA damage.

The observations outlined above suggested that Rtt107 cooperated with Rad55 to limit LOH and mitotic RCOs, and we therefore hypothesized that Rtt107 may limit genome instability through a DDR pathway that is known to be regulated by Rad55. Given that Rtt07 may not be required for Rad55’s canonical DSB repair function [20], we considered that Rtt107 may cooperate with Rad55 in regulating DNA damage tolerance, as mutants deficient in recombination can rely on tolerance to limit spontaneous LOH [59]. To test our hypothesis, we compared the spontaneous LOH phenotype of the *rtt107*Δ mutant with those of a *rev3*Δ mutant deficient in TLS, a *ubc13*Δ mutant deficient in error-free PRR, and a *rad18*Δ mutant deficient in both pathways [60,61]. Consistent with previous work, the *rad18*Δ mutant exhibited a modest (∼2-fold) but significant increase in spontaneous LOH [39] (Fig. S3). The *ubc13*Δ mutation did not affect the rate of spontaneous LOH in wild-type or *rtt107*Δ mutant, indicating that Rtt107 did not primarily limit LOH by promoting error-free PRR. In addition, the *rev3*Δ mutation increased the LOH phenotype of the *rtt107*Δ mutant, although this effect was modest (∼1.5-fold). The findings outlined above indicated that Rtt107 acted in the same pathway as Rad55 to limit spontaneous LOH and mitotic RCOs, but that Rtt107 did not cooperate with Rad55 in all contexts.

### 3.4 The *rad55-S404A* mutation largely disrupted the Rtt107–Rad55 interaction

To further delineate the mechanisms underlying the genetic relationship between *RTT107* and *RAD55*, we tested whether the physical interaction between Rtt107 and Rad55 was important for preventing genome instability. We considered that the Rtt107–Rad55 interaction may be mediated by Rad55-S404 phosphorylation, which does not require DNA damage induction [62], as the Rtt107–Rad55 interaction occurs under normal growth conditions [20]. To test this hypothesis, we generated a *rad55-S404A* mutation and established that it did not affect Rad55 protein level (Fig. S4). To determine whether the *rad55-S404A* mutation affected the Rtt107– Rad55 interaction, we performed a small-scale interaction assay using FLAG-Rad55 from log-phase cultures (Fig. 5A). We observed copurification of Rad57-HA and Rtt107-VSV with Rad55, which was consistent with a previous report [20]. The *rad57*Δ mutation did not eliminate the copurification of Rtt107 with Rad55, indicating that the Rtt107–Rad55 interaction was independent of Rad57. As Rtt107 was also found in Rad57 immunoprecipitations [20], these results were consistent with a model in which Rtt107 interacts with Rad55 to form an Rtt107– Rad55–Rad57 trimeric complex. Of note, the *rad55-S404A* mutation substantially reduced the amount of Rtt107, but not Rad57, that copurified with Rad55, suggesting that the Rad55-S404 residue was indeed important for the interaction with Rtt107, while having no appreciable effect on the Rad55–Rad57 interaction. For our purposes, we therefore concluded that this mutation was appropriate for investigating the importance of the Rtt107–Rad55 interaction in functional assays.

**Fig. 5.**
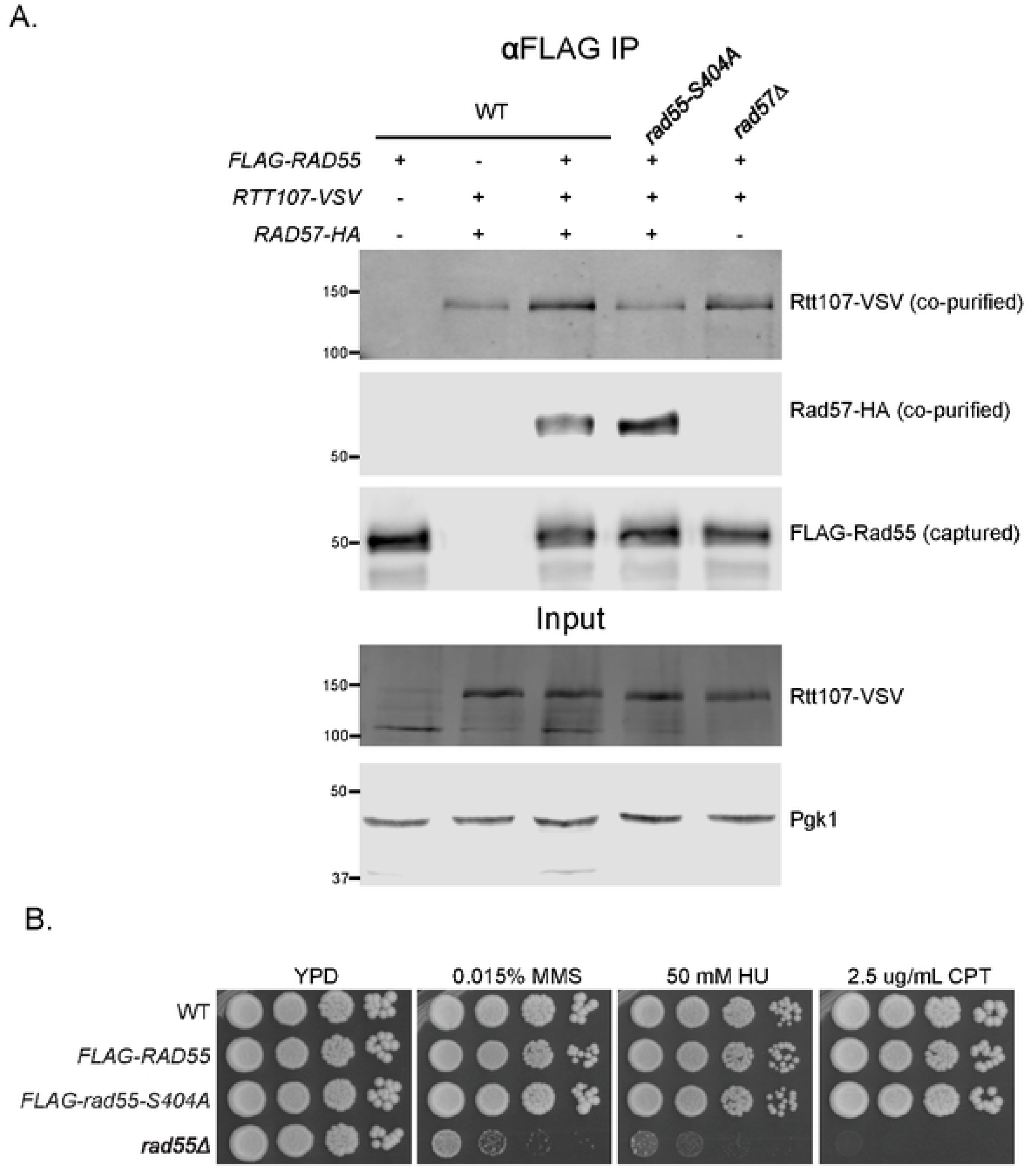
A *rad55-S404A* mutation largely disrupted the physical Rtt107–Rad55 interaction, but did not clearly affect growth during chronic exposure to DNA-damaging agents. Experiments were performed in the W303 *RAD5* background. A) The *rad55-S404A* mutation greatly reduced the amount of Rtt107 that co-purified with Rad55, but left Rad55–Rad57 interaction largely intact. Co-purification of Rtt107 with Rad55 did not require *RAD57.* FLAG purification was performed on log-phase cultures. Immunoblotting was performed using anti-FLAG, anti-VSV, and anti-HA antibodies. Anti-PGK1 antibody was applied to the input fraction as a loading control, due to the poor visibility of FLAG-Rad55 and Rad57-HA in whole cell extract. B) The *rad55-S404A* mutation or N-terminal 3×FLAG tag on Rad55 did not affect DNA damage sensitivity. Higher concentrations of DNA-damaging agents were used to examine the sensitivity of the *rad55-S404A* mutant, and the *rad55*Δ mutant was included for comparison. Tenfold serial dilutions of overnight cultures were plated on media containing the indicated concentrations of DNA-damaging agents, and then incubated for 2 days at 30°C.

A previous report suggested that mutation of the Rad55-S404 site does not affect the ability of cells to grow during chronic treatment with MMS [62]. As the phenotypes of DNA repair mutants can be highly dependent on the type of DNA-damaging agent used as well as the strain background, we examined whether the *rad55-S404A* mutation affected the ability of our strains to grow during chronic exposure to MMS, HU, or CPT (Fig. 5B). We found that the *rad55-S404A* mutation did not cause sensitivity to any of these agents at concentrations that severely inhibited growth of the *rad55*Δ mutant. Taken together, these observations suggested that the *rad55-S404A* mutation left Rad55 function in this context largely intact and that the interaction between Rtt107 and Rad55 was not crucial for growth during chronic exposure to DNA-damaging agents.

### 3.5 The *rad55-S404A* mutation increased spontaneous LOH and RCO rates

As the *rad55-S404A* mutation largely disrupted the physical Rtt107–Rad55 interaction, we next used it to examine whether the interaction between Rtt107 and Rad55 contributed to limiting spontaneous genome instability. First, we again employed the bimater assay to examine LOH induced by a broad range of events. The *rad55-S404A* mutation caused a modest (∼2-fold) increase in spontaneous LOH in comparison to the wild-type strain (Fig. 6A), consistent with a model in which Rtt107 and Rad55 prevented spontaneous RCOs in part by binding to each other. However, we noted that the *rad55*Δ mutation caused a much greater increase in LOH rate in the bimater assay (∼10-fold) (Fig. 4A). Further, the LOH phenotype of the *rtt107*Δ mutant was also greater than that of the *rad55-S404A* point mutant, and the *rtt107*Δ *rad55-S404A* double mutant exhibited a higher rate of LOH than either single mutant alone, suggesting that the function of Rtt107 in preventing LOH may not be solely dependent on the Rad55-S404 site and the Rtt107– Rad55 interaction To more granularly determine the types of spontaneous genome instability that may be prevented by Rtt107–Rad55 interaction, we again used the sectoring assay to specifically examine mitotic RCO events. The *rad55-S404A* mutation increased the frequency of mitotic RCOs, indicating that the Rad55-S404 site contributed to the limitation of spontaneous RCOs, and suggesting that the interaction between Rtt107 and Rad55 may be important in this context (Fig. 6B). The *rtt107*Δ *rad55-S404A* double mutant phenotype was similar to that of the *rtt107*Δ single mutant, and this pattern was broadly similar to the epistasis observed between the *rtt107*Δ and *rad55*Δ mutations in the context of the RCO phenotype (Fig. 4B), consistent with a shared role for Rtt107 and the Rad55-S404 site in limiting mitotic RCOs.

**Fig. 6.**
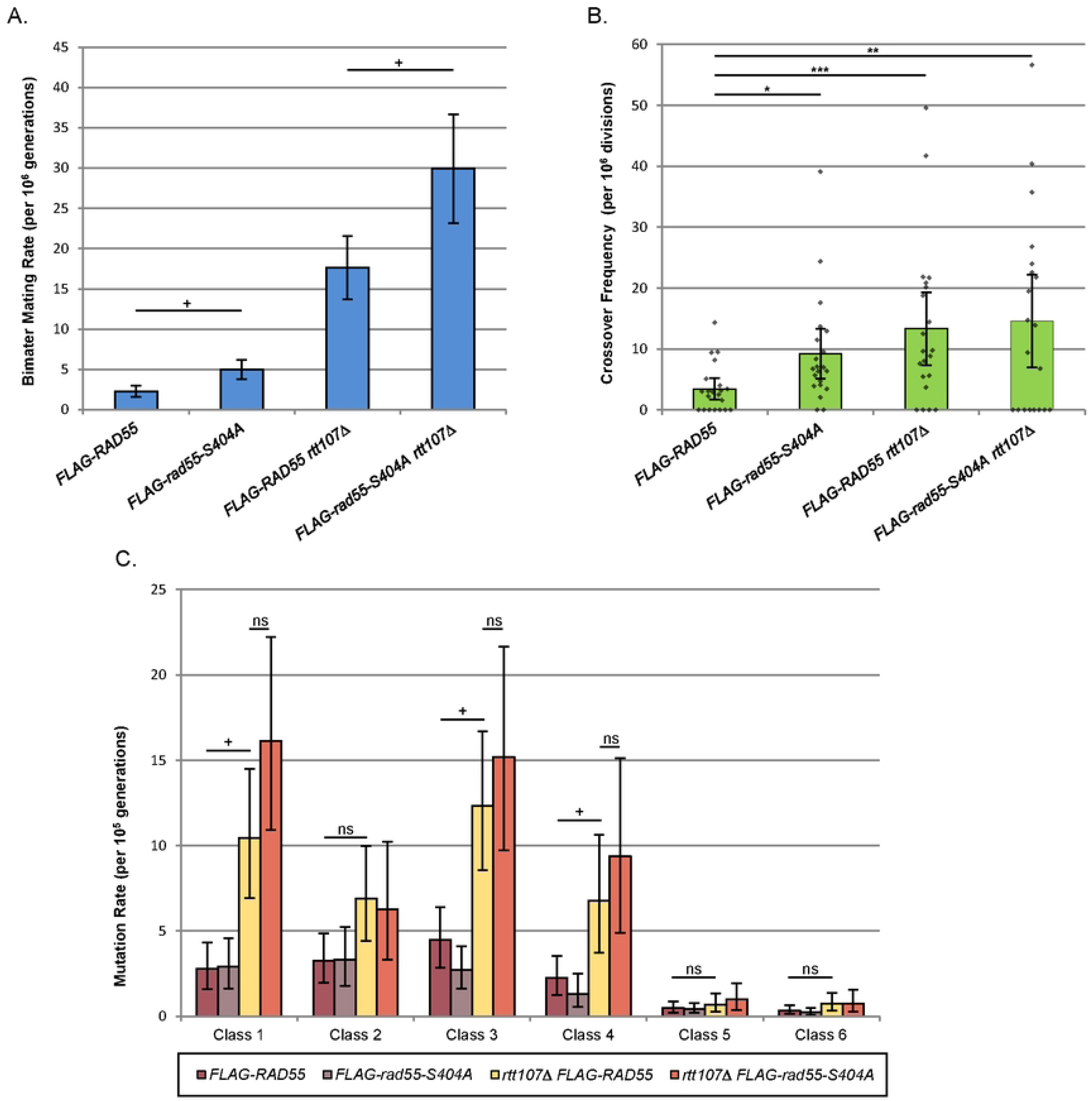
The *rad55-S404A* mutation increased spontaneous rates of LOH and mitotic RCOs. Experiments were performed in the W303 *RAD5* background. A) The *rad55-S404A* mutation caused a higher rate of LOH in the bimater assay, but did not exhibit epistasis with the *rtt107*Δ mutation. Mating rates were calculated from 10–12 independent colonies as described for the bimater assay in Figure 2B. +, non-overlap of 95% confidence intervals between two strains (statistically significant); ns, overlap of confidence intervals (not significant). B) The *rad55-S404A* mutation increased the frequency of mitotic RCOs, and the *rad55-S404A rtt107*Δ double mutant phenotype was similar to that of the *rtt107*Δ mutant. Frequencies were calculated from 20–21 independent colonies. *, *p* < 0.05; **, *p* < 0.01, ***, *p* < 0.005 compared to the wild-type (Student’s *t* test). C) The *rtt107*Δ mutant exhibited multiple types of spontaneous genome instability, and these phenotypes were not significantly affected by the *rad55-S404A* mutation. Six phenotypic classes of colony were scored using an assay for several types of mutation and mitotic recombination events. Rates were calculated and compared as described for the bimater assay in Figure 2B. +, non-overlap of 95% confidence intervals between two strains (statistically significant); ns, overlap of confidence intervals (not significant). Eight to nine independent colonies were tested for Class 1–4 events, and five to six colonies were tested for Class 5/6 events.

To further delineate the functional relevance of the Rtt107–Rad55 interaction, we considered that the spontaneous LOH and RCO phenotypes of the *rad55-S404A* mutant may be indicative of a defect in a specific DNA repair pathway, possibly the crossover-generating homologous recombination pathway. We therefore examined whether the *rad55-S404A* and *rtt107*Δ mutants exhibited various types of spontaneous genome instability related to recombinational repair using an established assay that discriminates between several distinct types of recombination and mutation in diploid cells (Figure 1) (see [49] for details). In this assay, Class 1/2 events primarily represent RCOs and break-induced replication (BIR) events. Class 3/4 events primarily represent local gene conversions that occur without crossovers, or *de novo* mutations occurring within the markers. Class 5/6 events primarily represent loss of one copy of chromosome V by nondisjunction. Using this approach, we found that the *rtt107*Δ mutant exhibited higher rates of Class 1 and Class 3/4 events but wild-type rates of Class 5/6 events (Fig. 6C). In the case of Class 2 events, the *rtt107*Δ mutant showed a trend toward a higher spontaneous mutation rate than wild-type, but this increase was not statistically significant. We further examined the Class 3 events in the *rtt107*Δ mutant by colony PCR and found that 24 of 28 events (86%) represented gene conversion rather than *de novo* mutation within the marker (see *Materials and Methods*). These observations suggested that *rtt107*Δ mutants have elevated rates of gene conversion, RCOs, and possibly BIR, but not chromosome loss. The *rad55-S404A* single mutant did not exhibit significantly increased rates in any of these classes, and the *rtt107*Δ *rad55-S404A* double mutant resembled the *rtt107*Δ single mutant. In contrast to the importance of the Rad55-S404 phosphorylation site for limiting LOH and RCOs, these results indicated that Rad55-S404 was not essential to limit multiple other types of spontaneous genome instability that are common in *rtt107*Δ mutants.

### 3.6 *RTT107* and *RAD55/RAD57* acted in similar genetic pathways under normal growth conditions

Our findings suggested that Rtt107 cooperated with Rad55 to limit spontaneous LOH and RCOs, and that this function was at least partially distinct from its role in the response to chronic DNA-damaging agent exposure. Therefore, we hypothesized that the genetic interaction spectrum of *RTT107* drawn from studies that did not apply DNA-damaging agents may provide relevant insights regarding the relations between *RTT107* and *RAD55*, and potentially also *RAD57*. To test this hypothesis, we compared published genetic interaction data on *RTT107*, *RAD55*, and *RAD57* from several high-throughput screens performed under normal growth conditions, and determined which genetic interactions were shared between at least two of these genes (Table 1) [63–71]. Of the 99 genetic interactions shared between *RAD55* and *RAD57*, we found that 40 were also shared with *RTT107.* This represented significant overlap between genes that interacted with *RTT107* and genes that interacted with *RAD55* + *RAD57* (Fisher’s exact test, two sided, *p* < 2.2 × 10^−16^). Fewer interactions (8) were shared between either *RTT107* + *RAD55* or *RTT107* + *RAD57* only. While further analyses will be required to fully establish the relationships between *RTT107*, *RAD55*, and *RAD57*, the greater than expected by chance overlap of shared genetic interactors suggested that these genes may act, at least in part, in the same pathways under normal growth conditions.

**Table 1.**
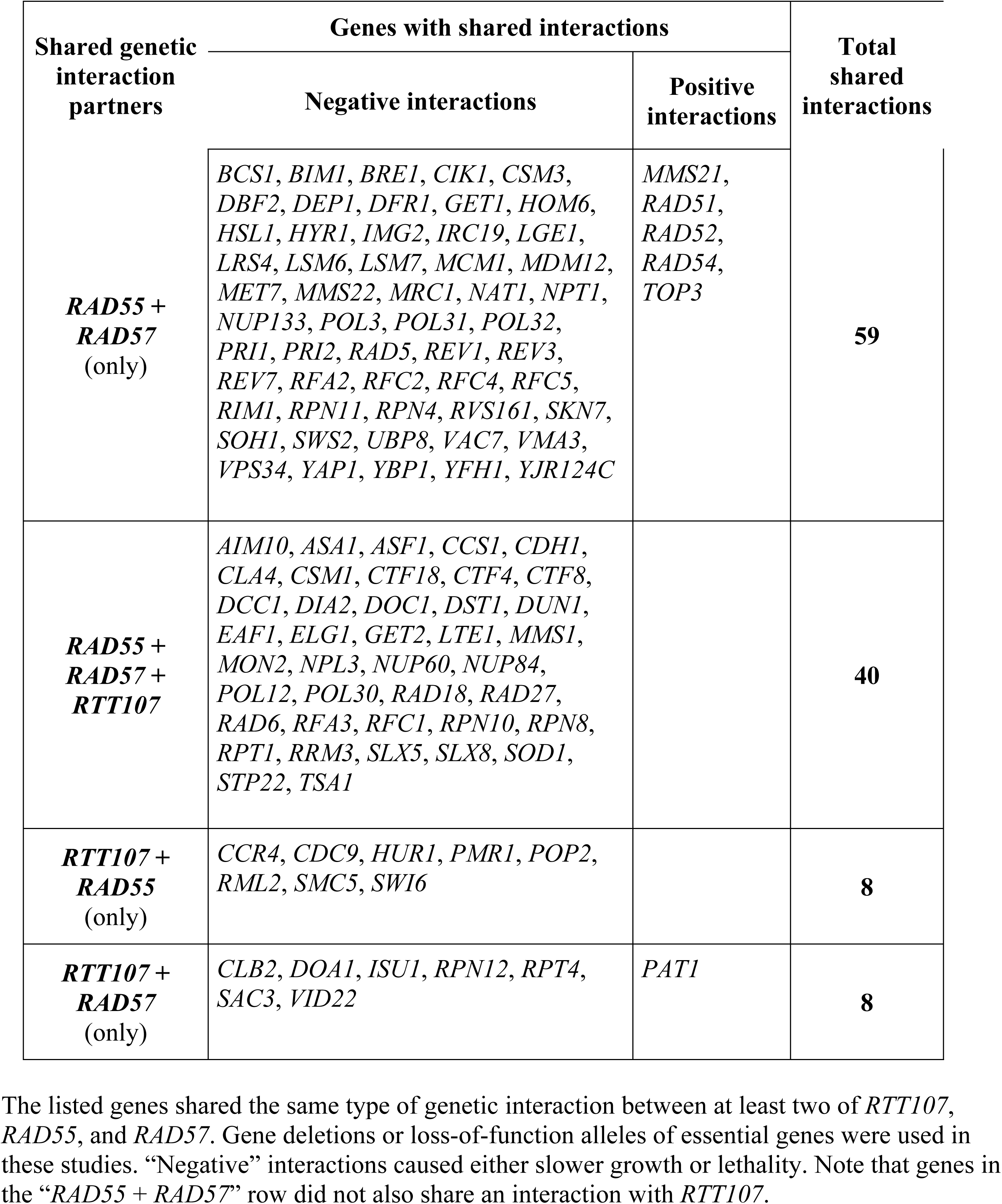
Shared and distinct genetic interaction profiles between *RTT107*, *RAD55,* and *RAD57*.

## 4 Discussion

Although Rtt107 is known as a multi-BRCT scaffold with homologs in fission yeast and vertebrates [11], its specific yet important function in limiting spontaneous genome instability phenotypes has remained unclear. Here, we showed that the role of Rtt107 in preventing spontaneous LOH was distinct from its well-characterized functions as a scaffold that localizes to γH2A after induction of DNA damage. Instead, Rtt107 cooperated with the recombination mediator Rad55, at least in part, to limit spontaneous LOH and mitotic RCOs in diploid cells.

The Rad55-S404 phosphorylation site was partially required for the Rtt107–Rad55 protein– protein interaction and to limit spontaneous LOH and RCOs, suggesting that the physical interaction between Rtt107 and Rad55 contributed to the suppression of these events. In contrast, the cooperation between Rtt107 and Rad55 did not extend to prevention of all types of spontaneous genome instability. For example, Rtt107 acted in the same pathway as Slx4 to limit rDNA recombination. Taken together, our results indicated that Rtt107 acted with either the recombination mediator Rad55 or its well-characterized binding partner Slx4 to limit types of genome instability that occur spontaneously under normal growth conditions.

### 4.1 Rtt107 limited spontaneous genome instability in distinct pathways

The role of Rtt107 in limiting genome instability differed between various types of instability. Rtt107 cooperated with Rad55 to prevent LOH at the mating locus and mitotic RCOs on chromosome V in diploid cells, likely in part by binding to Rad55 in a manner dependent on Rad55-S404 (Fig. 7A). Our data were also consistent with a model in which Rtt107 interacts with the complete Rad55–Rad57 heterodimer by binding primarily to Rad55, and genetic interaction data suggested that *RTT107*, *RAD55*, and *RAD57* can act in similar pathways under normal growth conditions (Table 1). However, the Rad55-S404 site, while important for mediating the physical interaction between Rad55 and Rtt107, was not needed to limit other types of instability that are common in diploid cells, specifically gene conversion and possibly BIR (Fig. 6C). These observations suggested that Rtt107 primarily acted independently of the Rtt107–Rad55 interaction to limit these events, whereas the Rtt107–Rad55 interaction may specifically limit LOH and mitotic RCOs. We also found that the function of Rtt107 in the context of LOH was dependent on BRCT5/6 but independent of the Rtt107-K887 residue that mediates interaction with γH2A (Fig. 2B) [10,19]. This was in contrast to earlier observations indicating that *rtt107-K887M* and *rtt107*Δ*BRCT5/6* mutants exhibit similar phenotypes with respect to growth during chronic MMS treatment and the recruitment of Rtt107 to induced DSBs [10]. Interestingly, the structure of BRCT5/6 is remarkably flexible, and protein binding via BRCT5/6 using surfaces other than the γH2A-binding pocket has been proposed previously [19]. Taken together, these findings suggested that Rtt107–BRCT5/6 may bind to a non-γH2A partner and thus prevent LOH (Fig. 7A), and the identification of such a partner represents an opportunity for future research. It is of interest to consider that Rad55 binds to Rtt107–BRCT1–4 [14,20], and that Rtt107 may act to bridge Rad55, or the complete Rad55–Rad57 heterodimer, to a partner that binds BRCT5/6.

**Fig. 7.**
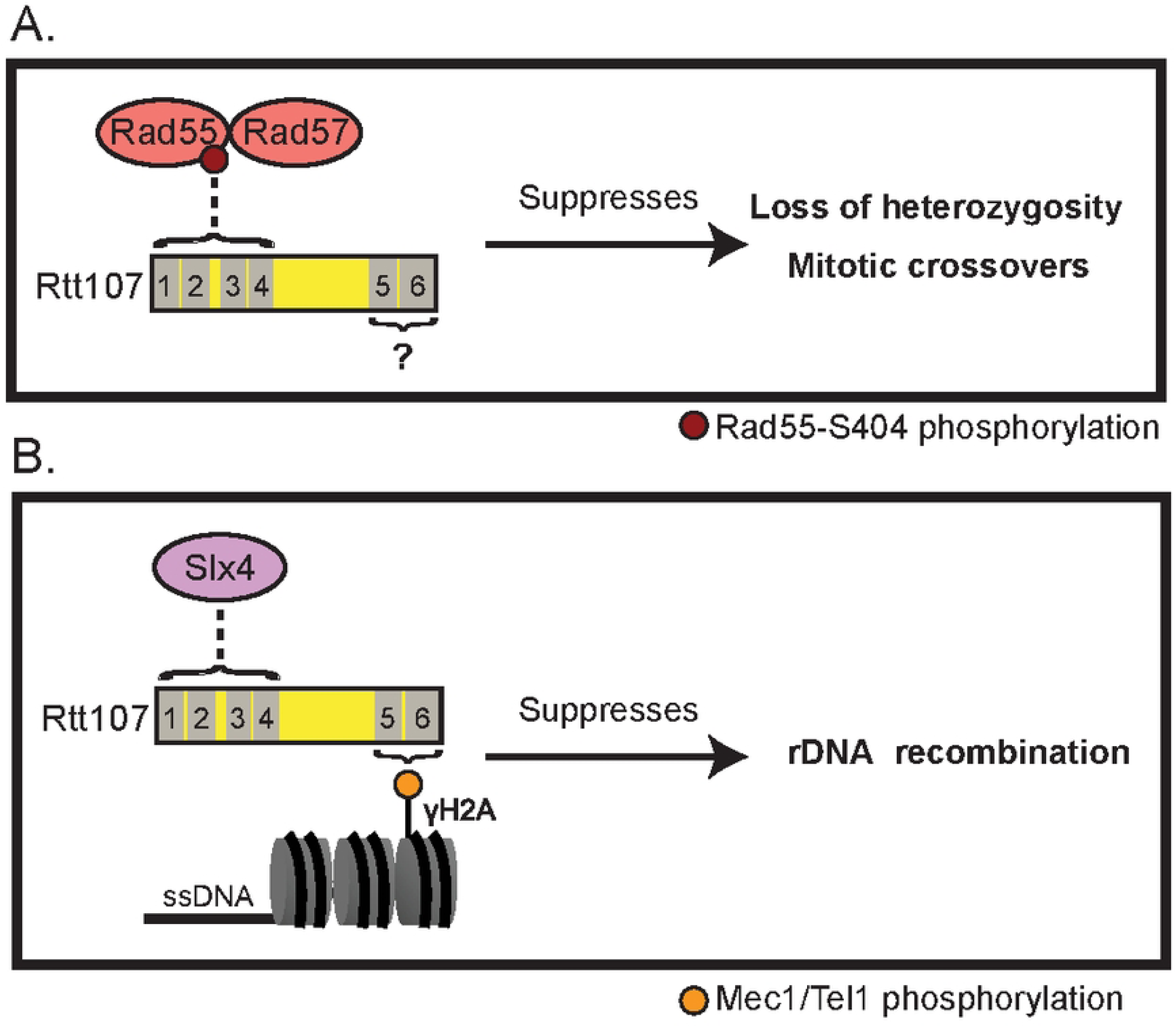
Rtt107 acts with different partners to prevent various types of genome instability. The dotted lines indicate physical interactions between Rtt107 and proteins that maintain stability in the same pathway. Numbered gray boxes indicate Rtt107 BRCT domains. A) Rtt107 prevents spontaneous LOH and mitotic RCOs by cooperating with the Rad55–Rad57 heterodimer, which binds to Rtt107–BRCT1–4. The Rad55-S404 phosphorylation site contributes to the Rtt107–Rad55 interaction. Rtt107–BRCT5/6 contributes to the function of Rtt107 in preventing LOH events, possibly by binding to a partner other than γH2A. B) Rtt107 prevents rDNA recombination by cooperating with Slx4, which binds to Rtt107–BRCT1–4. Rtt107 may act as a γH2A-binding scaffold that recruits Slx4 near lesions in this context.

Rtt107 cooperated with Slx4 to prevent hyperrecombination at the rDNA locus in haploid cells, while not depending primarily on Rad55 in this context. Recent work suggested that Rtt107 limits rDNA recombination at least in part as a γH2A-binding scaffold for its physical interaction partners, including Slx4 [14]. Therefore, the cooperation of Rtt107 with Slx4 in rDNA maintenance may involve binding to γH2A via Rtt107–BRCT5/6 (Fig. 7B). We note that Rtt107 did not rely on Slx4 to prevent spontaneous LOH (Fig. 3A), again suggesting that Rtt107 acts in different pathways to limit different types of spontaneous genome instability.

In addition to the spontaneous genome instability phenotypes examined in this study, the *rtt107*Δ mutant exhibits elevated levels of spontaneous DNA damage foci in S/G_2_/M phase [15,25,38]. Interestingly, the *rtt107*Δ mutation does not exhibit broad epistasis with mutations of canonical DNA damage repair or tolerance pathways [8,20,44,72,73]. Taken together, these findings were consistent with a model in which Rtt107 maintains genome stability at least in part by limiting the levels of DNA damage that occur during replication, rather than by directly affecting a specific repair process or pathway choice. Furthermore, it is interesting that DSBs appear to be the primary initiator of spontaneous crossovers in yeast [74–76]. The dispensability of Rtt107 for survival after exposure to DSB-inducing ionizing radiation [8], along with the requirement for Rtt107 in preventing spontaneous crossovers, could be reconciled by a model in which Rtt107 prevents the occurrence of DSBs but is not essential for DSB repair.

### 4.2 Cooperation between Rtt107 and Rad55 in genome maintenance

Our genetic evidence suggested that Rtt107 interacted with Rad55 to limit rates of spontaneous RCOs, and functioned at least in part with Rad55 to limit LOH. We note that the LOH phenotype examined in this study could have been largely driven by mitotic RCO events, and that these phenotypes would be consistent with a relationship between the Rtt107–Rad55 interaction and recombinational repair pathways, which are mediated by Rad55–Rad57.

However, it remains unclear how Rtt107 and Rad55 would cooperate to prevent spontaneous genome instability. In complexes such as Rtt107–Slx4, Rtt107–SMC5/6, or Rtt107–Rtt101– Mms1–Mms22, Rtt107 has commonly been proposed to act as a scaffold that recruits its binding partners to γH2A near DNA damage lesions (Fig. 7B) [9,10,14,16,55]. In contrast, our results presented here suggested that Rtt107–γH2A binding was actually dispensable for the function of Rtt107 in preventing LOH, and a model in which Rtt107 simply recruits Rad55 to lesions thus appears less applicable. Rtt107 and Rad55 also localize to different DNA damage signals, as Rad55 associates with the Rad51–ssDNA filament during recombination [26,77], while Rad51 does not appear to assemble nonspecifically onto double-stranded DNA [78]. Rtt107 localizes to the histone marks γH2A and H4T80ph, which flank DSB sites and are excluded from ssDNA [7,10,78]. Therefore, while Rtt107 and Rad55 both localize near DSBs in broad terms, their known recruitment pathways appear to target different regions around DSBs. Rad51 can be detected in similar regions to γH2A due to homology search performed by the Rad51 filament [78], but it is unclear whether the Rtt107–Rad55 interaction is important during homology search or some other process. Alternatively, Rtt107 and Rad55 may cooperate at sites other than DSBs, such as stalled replication forks. The localization of Rtt107 behind stressed replication forks has been shown to be γH2A-dependent [13], whereas the recruitment of Rad55 to replication forks is less well understood [79,80] and it is unclear whether Rtt107 and Rad55 can be recruited to stressed forks at similar times.

While our findings were consistent with a relationship between the Rtt107–Rad55 interaction and recombinational repair, it is also important to note that Rtt107 does not suppress recombination at all loci. For example, a genome-wide study that identified rDNA hyperrecombination in the *rtt107*Δ mutant found normal rates of recombination at the *LEU2* locus in two assays [38].

We found that the Rad55-S404 phosphorylation site was important for the physical interaction between Rtt107–Rad55 and for limiting mitotic RCOs and LOH. However, in the context of the LOH phenotype, we did not identify clear epistasis between the *rtt107*Δ and *rad55-S404A* mutations (Fig. 6A), which suggested that the Rad55-S404 site may have roles in Rad55’s function beyond the complexes or pathways involving Rtt107. Further, it is possible that residual binding between Rtt107 and Rad55 occurs in the *rad55-S404A* mutant, limiting the impact of the targeted *rad55-S404A* mutation on certain genome instability phenotypes, although this was not detectable in the assays employed here. In contrast, we observed epistasis between the *rtt107*Δ and *rad55*Δ mutations in the context of the LOH phenotype (Fig. 3F and 4A), suggesting that Rtt107 and Rad55 act broadly in the same pathway to prevent LOH.

It should also be noted that we identified cooperation between Rtt107 and Rad55 in assays using diploid yeast, whereas Rtt107 did not cooperate primarily with Rad55 to prevent rDNA recombination in haploids (Fig. 4D). Diploid and haploid yeast differ in DDR regulation, with diploids utilizing HR over NHEJ for DSB repair [30]. Further, the sensitivity of *rad55Δ* mutants to ionizing radiation is suppressed by heterozygosity at the mating locus, which occurs naturally in diploids [30,31]. Therefore, the degree of cooperation between Rtt107 and Rad55 in genome maintenance may be partly dependent on ploidy.

We identified a spontaneous LOH phenotype for an *rtt107-K426M* mutant (Fig. 2B), suggesting that the K426-dependent interaction of Rtt107 with phosphorylated H4T80 may also be important for preventing spontaneous LOH, possibly by affecting the timing of Rtt107 recruitment [7]. However, we note that K426 is within Rtt107–BRCT1–4, and that BRCT1–4 interacts with a number of interaction partners of Rtt107, including Rad55 [9,14,15,20]. Therefore, it remains possible that the *rtt107-K426M* mutation could also affect the Rtt107– Rad55 complex, or other Rtt107 complexes.

### 4.3 Cooperation between Rtt107 and Slx4 in rDNA maintenance

We found that Rtt107 acted in the same pathway as Slx4 to maintain rDNA stability (Fig. S2). The role of Rtt107 in rDNA maintenance may be similar to its role in GCR suppression in haploid cells, in which Rtt107 acts at least in part as a γH2A-binding scaffold for Slx4, Mms22, and Nse6 (Fig. 7B) [9,14,15,20]. It is not yet clear whether Rtt107 also cooperates with the Slx1 endonuclease in this context. Although an Rtt107–Slx1–Slx4 complex has been identified [15], Rtt107 has not been previously implicated in the endonuclease function of Slx1–Slx4. In addition, we note that the rDNA hyperrecombination phenotype of the *slx4*Δ mutant identified in this study differs from a previous report in which the *slx4*Δ mutant exhibited lower rates of rDNA recombination than wild-type [81]. This discrepancy may be related to measurement of the loss of a single *ADE2* marker in the rDNA in the earlier study, rather than the combined *ADE2-CAN1* marker used here and in a recent study identifying rDNA hyperrecombination in an *slx4* mutant [14,50]. Relatedly, the *Schizosaccharomyces pombe* homolog of Rtt107, Brc1, has been proposed to play a role in genome maintenance in regions that are susceptible to replication stress, including the rDNA [82], likely at least in part due to recruitment of Brc1 to γH2A, with the γH2A–Brc1 foci colocalizing with rDNA [83].

Recent work identified GCR and rDNA recombination phenotypes for mutations of the Rtt107 interaction motifs (RIMs) that are important for the interaction between Rtt107–BRCT1– 4 and three of the binding partners of Rtt107, i.e., Slx4, Mms22, and the SMC5/6 subunit, Nse6 [14]. These *RIM* mutants each exhibited mild rDNA hyperrecombination phenotypes in comparison to the more pronounced phenotype of the *rtt107*Δ mutant, suggesting that individual interactions may have had limited or partially redundant contributions to the rDNA maintenance function of Rtt107. Our results were consistent with substantial cooperation between Rtt107 and Slx4 in rDNA maintenance, as the *slx4*Δ and *rtt107*Δ mutations had similar and epistatic effects on rDNA stability (Fig. S2).

### 4.4 Distinct roles for Rtt107 in genome maintenance and DNA damage resistance

In this study, we found that the *rad9-2A* and *H2A-S129A* mutations increased the LOH rate of the *rtt107*Δ mutant (Fig. S1 and 2C), despite suppressing the hypersensitivity of the *rtt107*Δ mutant to chronic MMS treatment [16,17,22]. Previously, we reported a similar pattern in which the *dot1*Δ mutation suppressed the MMS sensitivity of the *rtt107*Δ mutant, but did not suppress its spontaneous LOH phenotype [25]. Each of these mutations that suppress the MMS sensitivity of the *rtt107*Δ mutant can also dampen DNA damage checkpoint activity [53,84–87], and the *H2A-S129A* mutation additionally suppresses the MMS-induced checkpoint hyperactivation phenotype of the *rtt107*Δ mutant [16,17]. The lack of suppression by these mutations of the LOH phenotype in the *rtt107*Δ mutant reported here suggested that the role of Rtt107 in preventing spontaneous LOH was largely distinct from its role in preventing treatment-induced checkpoint hyperactivation. This distinction was further emphasized by the observations that the *rtt107*Δ and *rad55*Δ mutations exhibited epistasis in the context of the LOH phenotype (Fig. 3F and 4A) and a negative genetic interaction in the context of growth during MMS exposure (Fig. 4E) [20]. We also found that *rtt107*Δ mutation suppressed the LOH phenotype of the *mms22*Δ mutant (Fig. 3C), despite the additive effects of *rtt107*Δ and *rtt101*Δ mutations on MMS sensitivity observed in a growth assay [88]. Taken together, these findings indicated that while abundant information is available on the genetic interactions of the *rtt107*Δ mutant in the context of exposure to DNA-damaging agents, these data may not directly reflect the function of Rtt107 in preventing spontaneous LOH.

The lack of a DNA damage sensitivity phenotype for the *rad55-S404A* mutant suggested that the Rtt107–Rad55 interaction was not important for DNA damage resistance (Fig. 5B). This was unsurprising, as the *rtt107*Δ mutant was less sensitive to DNA-damaging agents than the *rad55*Δ mutant (Fig. 4E). In addition, a previous genome-wide screen in a different genetic background performed in our laboratory revealed a negative genetic interaction between the *rtt107*Δ and *rad55*Δ mutations in the context of CPT sensitivity [73]. However, there was no clear negative interaction between the *rtt107*Δ and *rad55*Δ mutations at this CPT concentration in the W303 *RAD5* genetic background, although the sensitivity of the *rtt107*Δ *rad55*Δ double mutant was slightly greater than that of the *rad55*Δ single mutant (Fig. 4E).

Taken together, the findings presented here showed that Rtt107 plays multiple roles in the prevention of spontaneous genome instability, which are often distinct from its previously studied functions in the response to DNA-damaging treatments. Further studies are required to determine the specific mechanisms by which Rtt107 prevents various types of genome instability with its diverse interaction partners, including Rad55 and Slx4.

## Supporting information

Supplementary raw data S1

## Acknowledgments

We thank G. Leung for preliminary work in strain construction; W. D. Heyer, J. Diffley, C. D. Allis, B. Pfander, L. Aragon, and J. Krebs for generously providing yeast strains and plasmids; and M. Aristizabal, P. Stirling, W. D. Heyer, and A. Kerr for critical reading of the manuscript and suggestions.

## Supplementary Figure Captions

**Fig. S1. Rtt107 did not limit spontaneous LOH by regulating the Rad9–Dpb11 interaction.**

The *rad9-2A* allele enhanced the spontaneous LOH rate of the *rtt107*Δ mutant in the bimater assay. Mating rates were calculated and compared as described for the bimater assay in Figure 2B. +, non-overlap of 95% confidence intervals between two strains (statistically significant); ns, overlap of confidence intervals (not significant). This experiment was performed in the W303 *rad5-535* background, and 12 independent colonies were tested.

**Fig. S2. Rtt107 limited rDNA recombination in the same pathway as Slx4.**

The *rtt107*Δ, *slx4*Δ, and *rtt107*Δ *slx4*Δ mutants exhibited similar rDNA hyperrecombination phenotypes. rDNA recombination rates and 95% confidence intervals were determined as described in Figure 4D. +, non-overlap of 95% confidence intervals between two strains (statistically significant); ns, overlap of confidence intervals (not significant). This experiment was performed in the W303 *RAD5* background, and eight independent colonies were tested.

**Fig. S3. Rtt107 did not primarily limit spontaneous LOH by regulating DNA damage tolerance pathways.**

Deletion of *UBC13* did not affect the rate of LOH in the wild-type or *rtt107*Δ mutant. Mating rates were calculated and compared as described for the bimater assay in Figure 2B. +, non-overlap of 95% confidence intervals between two strains (statistically significant); ns, overlap of confidence intervals (not significant). This experiment was performed in the W303 *rad5-535* background, and 11-12 independent colonies were tested.

**Fig. S4. The *rad55-S404A* mutation did not affect Rad55 protein level.**

Immunoblotting was performed using anti-FLAG antibody on whole-cell extract. Anti-PGK1 antibody was used as a loading control.

