## Supplementary figures and images for "Rtt107 cooperates with Rad55 or Slx4 to maintain genome stability in *Saccharomyces cerevisiae*"

### Supplementary Figures

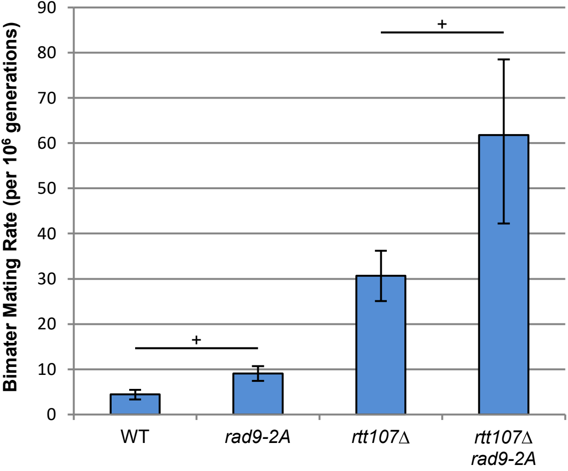

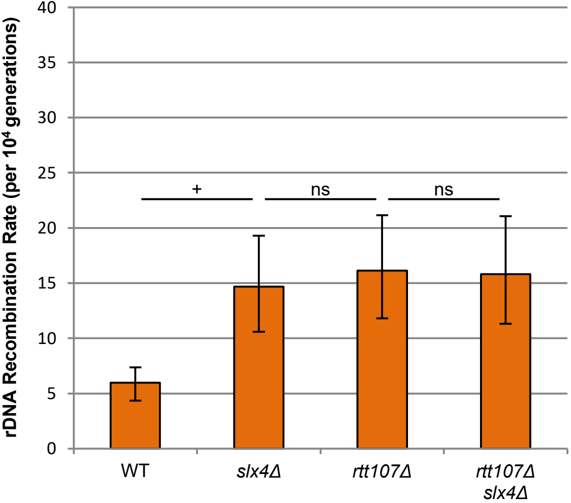

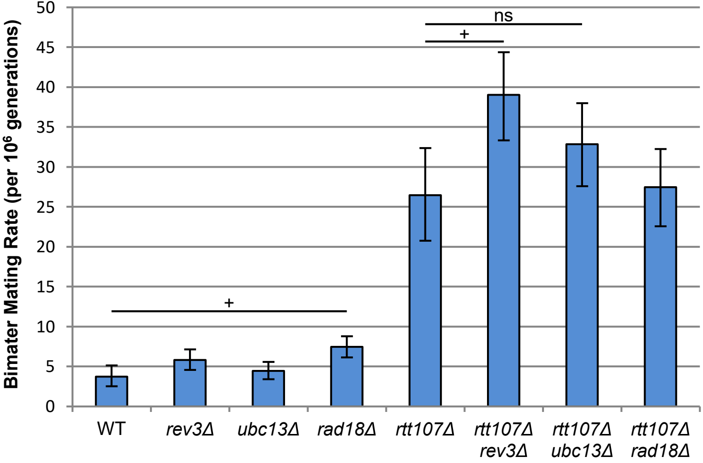

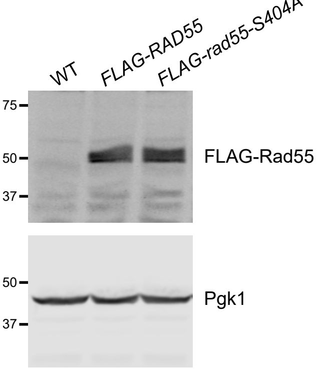


**Fig. S4.**
